# Pervasive sublethal effects of agrochemicals as contributing factors to insect decline

**DOI:** 10.1101/2024.01.12.575373

**Authors:** Lautaro Gandara, Richard Jacoby, François Laurent, Matteo Spatuzzi, Nikolaos Vlachopoulos, Noa O Borst, Gülina Ekmen, Clement M Potel, Martin Garrido-Rodriguez, Antonia L Böhmert, Natalia Misunou, Bartosz J Bartmanski, Xueying C Li, Dominik Kutra, Jean-Karim Hériché, Christian Tischer, Maria Zimmermann-Kogadeeva, Victoria Ingham, Mikhail M Savitski, Jean-Baptiste Masson, Michael Zimmermann, Justin Crocker

## Abstract

Insect biomass is declining across the globe at an alarming rate. Climate change and the widespread use of pesticides have been hypothesized as two underlying drivers. However, the lack of systematic experimental studies across chemicals and species limits our causal understanding of this problem. Here, we employed a chemical library encompassing 1024 different molecules—including insecticides, herbicides, fungicides, and plant growth inhibitors —to investigate how insect populations are affected by varying concentrations of pesticides, focusing on sublethal doses. Using a controlled laboratory pipeline for *Drosophila melanogaster*, we found that 57% of these chemicals affect the behavior of larvae at sublethal concentrations, and an even higher proportion compromises long-term survivability after acute exposure. Consistent with these results, we observed that exposure to chemicals at doses orders of magnitude below lethality induced widespread phosphorylation changes across the larval proteome. The effects of agrochemicals were amplified when the ambient temperature was increased by four degrees. We also tested the synergistic effects of multiple chemicals at doses found widely in nature and observed fitness-reducing changes in larval developmental time, behavior, and reproduction. Finally, we expanded our investigation to additional fly species, mosquitos, and butterflies and detected similar behavioral alterations triggered by pesticides at sublethal concentrations. Our results provide experimental evidence that strongly suggests sublethal doses of agrochemicals coupled with changes in environmental temperatures are contributing to the global decline in insect populations. We anticipate that our assays can contribute to improving chemical safety assessment, better protect the environment, secure food supplies, and safeguard animal and human health, as well as understand our rapidly changing world.

## Main

During the past decade, numerous reports have documented global declines in insects and their biodiversity^1–3^. Changes in species diversity and high rates of compositional turnover have been reported worldwide^4,5^. Field studies have identified several factors associated with this decline, including habitat loss due to extensive farming and urbanization, global warming, and the widespread use of pesticides^1–3^. However, these factors are rarely causally attributed to biodiversity trends.

Sublethal concentrations of pesticides have been proposed to be a major driver of the loss of insect biodiversity^2^. Case studies have outlined how sublethal doses of such agricultural chemicals can alter multiple aspects of insect biology, including metabolism, development, immunology, fecundity, and behavior^6^. Thus, even when employing pesticides with low non-target activity (i.e., LD50_pest spp._<<LD50_non-pest spp._), side effects on non-pest insect species may still occur, as sublethal impacts are generally not accounted for in these tests^7^. However, a lack of systematic experimental studies limits our mechanistic understanding of the scale and magnitude of this problem.

Here, to explore the sublethal effects of agrochemicals on *Drosophila melanogaster*—an insect model system for toxicology studies^8^—we established a quantitative analysis platform to assess the effects of these molecules on larval behavior, physiology and fitness.

### Behavior screen using an agrochemical library

We focused on the larval stage to make compound delivery precise^8^ and to increase the throughput of the assay. We studied the effects of exposing larval populations (∼25 3^rd^ instar larvae per replicate; three replicates per pesticide and concentration) across three concentrations (2 μM, 20 μM, and 200 μM) of 1024 different agrochemicals (Table S1) for 16 hours, in multiwell plates containing liquid food^9^ (Fig. 1a-c). Constant liquid exposure allows consistent delivery across experiments, mitigating the effects of toxin avoidance^10^. 200 and 20 μM are the concentrations used for field applications for many pesticides^11,12^—and thus, they constitute realistic concentrations for agricultural areas at the moment of spraying. 2 μM is a level where many of the chemicals can be detected in natural habitats. For example, previous work assessing the environmental occurrence of glyphosate across the US reported glyphosate concentrations as high as 2.5 μM in ditches and 1.8 μM in wetlands^13^. After this controlled incubation, we measured the number of larvae that survived acute exposure (Fig. 1c) while simultaneously capturing videos of them moving in a 2D arena to characterize the behavioral effects of these treated animals (behavioral shifts, Fig. 1c). The surviving larvae were then transferred to standard pesticide-free solid food (see Methods) and the number of adult flies emerging in those populations was measured after 10 days (long-term lethality, Fig. 1c).

**Fig.1:**
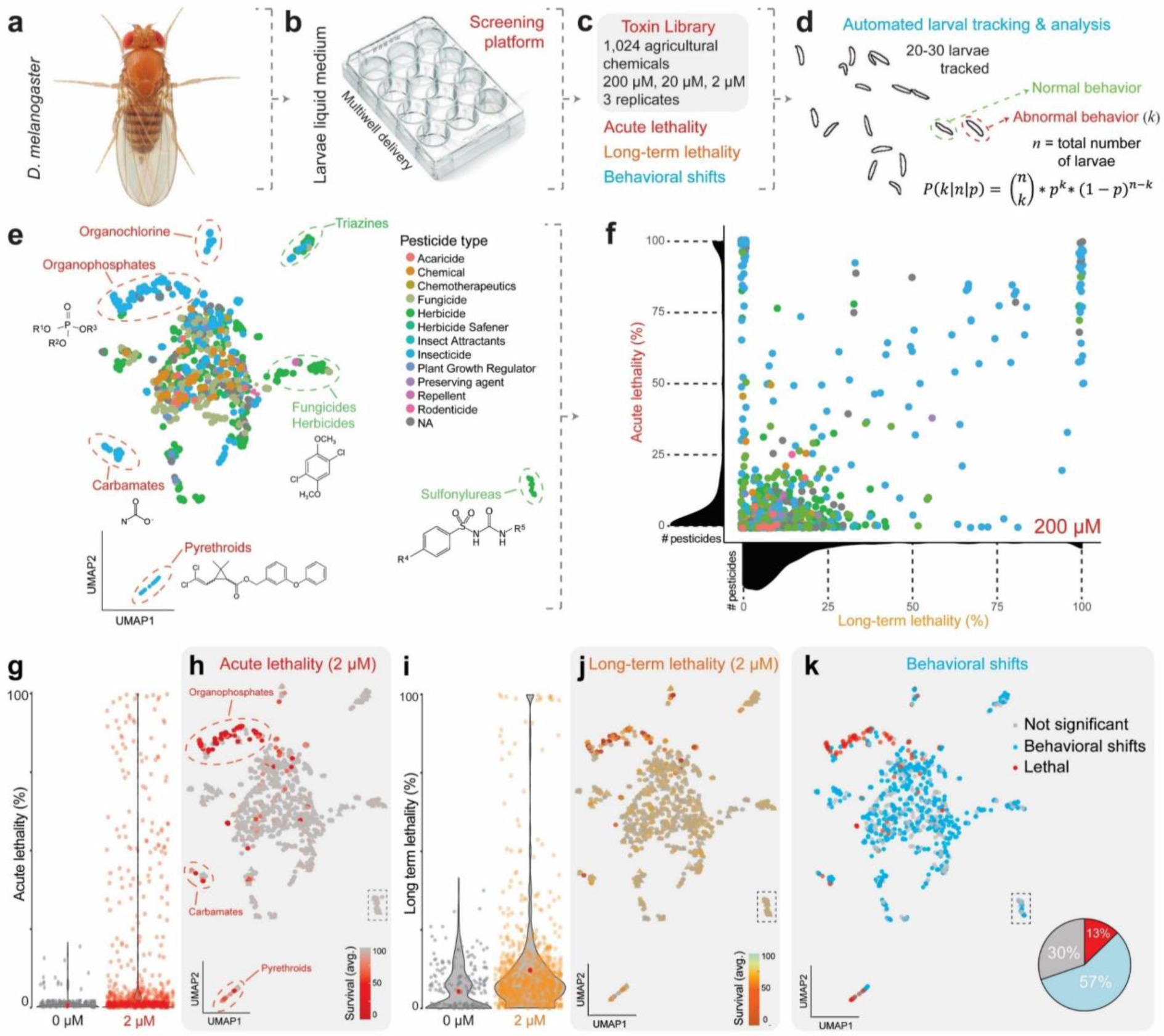
Agrochemicals alter larval development and behavior at sublethal concentrations. **a-d**, Schematic of the screen used for analyzing sublethal effects of pesticides. Third instar *D. melanogaster* larvae (**a**) were exposed overnight in liquid medium (**b**) to a broad agrochemical panel, at multiple concentrations (**c**), and (immediate) acute lethality, long-term lethality and the behavioral state of the treated populations were quantified after the exposure. The identification of aberrant behavior was based on the statistical comparison of the number of larvae showing altered frequencies of stereotypic behaviors in each treatment with control populations (**d**, see Methods). **e**, UMAP projections of the Morgan molecular fingerprints of the different molecules encompassed in the library, color-coded according to their pesticide type. **f**, Average (N=3, n∼20) acute and long-term lethality values observed for all the molecules in the library when employed at 200 μM. The color code represents the pesticide type, as in **e**. The axes also show the distribution of values for each of the variables. **g,** Average (N=3, n∼20) of acute lethality observed for all the tested pesticides when used at 2 μM. Red dots show the mean across all molecules. **h,** Acute lethality (as in **g**) mapped on top of the UMAP described in **e**. **i,** Average (N=3, n∼20) of long-term lethality observed for all the tested pesticides when used at 2 μM. Red dots show the mean across all molecules. **j,** long-term lethality (as in **i**) mapped on top of the UMAP described in **e**. **k**, Classification of pesticides either as lethal at 2 μM (red), significantly affecting behavior at sublethal concentrations (p<0.05 for at least the median number of replicates at that concentration, positive, light blue), or with no significant effect on behavior (negative, grey). This classification is shown as a color code on top of the UMAP described in **e**.

Automated analysis pipelines were established to extract behavioral information from the recorded videos (Fig. 1d). Individual larvae were tracked over time and their behavior was characterized either by extracting features from the trajectory followed by each larva or by classifying each larva at every frame of the video into seven different stereotypic behavioral classes using Larva-Tagger (see Methods). The impact of each treatment on larval behavior was tested at the population scale by comparing the features or classes distribution in each condition to the corresponding distribution in untreated controls using a standard outlier detection method (Fig. 1d, see Methods). Finally, data was imported to an open-access, relational database, including an interactive website (https://agrotoxin.embl.de) linked to the Image Data Explorer application^14^ (Extended Data Fig. 1a) for downstream applications.

### Sublethal effects of agrochemicals on insect physiology

To visualize the chemical diversity within our library, we applied the dimensionality reduction method UMAP^15^ to the Morgan molecular fingerprints of the compounds (Fig. 1e and Extended Data Fig. 1b). These fingerprints can be used for fast similarity comparisons that form the basis for structure–activity relationships. Notably, whilst some groups exhibited chemical identities that clustered based on the UMAP embedding (e.g., pyrethroid insecticides), there is a strong overlap in the chemical structure of most other groups (Extended Data Fig. 1b-d), which may help explain the potential effects of pesticides not formally classified as insecticides on insect systems^2^.

At the highest concentration tested, the majority of chemicals had limited acute or long-term toxicity (Fig 1f). Across the multiple doses tested, a general increase in acute lethality was observed at higher concentrations (Extended Data Fig. 2a). Each condition was tested in triplicate, with a high level of reproducibility at the level of acute lethality (Extended Data Fig. 2b). Long-term lethality had higher variability (Extended Data Fig. 2c), possibly due to the high initial lethality and resulting survivorship bias. Interestingly, acute and long-term lethality are not strongly correlated (Fig. 1f, R^2^=0.06 at 200 μM), as some pesticides with very strong acute lethality do not seem to affect *Drosophila* viability in the long term.

Unsurprisingly, substantial acute lethality at the lowest tested concentration was observed in chemicals classified as insecticides (i.e. organophosphates, pyrethroids, carbamates, etc; Fig. 1g-h). However, significant long-term lethality appears widespread across the library, with many non-insecticide pesticides influencing the survivability of the larval populations upon overnight exposure (Fig. 1i-j), even at the lowest tested concentration. A substantial fraction of the library (57%) reproducibly affected larval behavior (p<0.05, see Methods, Fig. 1k and Extended Data Fig. 1e), altering the frequencies of stereotypic behaviors (35%), trajectories (6%), or both (16%), even at sublethal concentrations (acute lethality < 10%)—these effects are not restricted to insecticides (Fig. 1k). Therefore, while some chemicals may not directly compromise viability in controlled laboratory environments, many cause behavioral changes (Extended Data Fig. 2d), which could compromise mating, feeding or other behaviors critical to survival^16,17^.

To cross-validate the sublethal effects of pesticides and to identify molecular signatures of pesticide exposure, we analyzed the phosphoproteome of larvae treated overnight with one of five highly prevalent pesticides (three chemically diverse insecticides, a fungicide, and a herbicide, Fig. 2a). Posttranslational modification of preexisting proteins is rapid and can occur before transcriptional responses to toxic stimuli^18^. Due to the high sensitivity of these assays, we exposed larvae at 0.2 μM—an even lower concentration but still within environmental levels. Even though these pesticides showed different degrees of acute lethality (Fig. 2b), all of them significantly affected larval behavior at this lower concentration (Fig. 2c). For example, glyphosate, a widely used herbicide across the world^19^ which is not lethal at 0.2 μM (Fig 2b), led to a significant increase in the frequency of turns (headcasting), paralleled by a decreased frequency of stops—a behavioral effect that was reflected in the trajectories of the larvae (Fig. 2d-e, Extended Data Fig. 3a-b). Based on the proteomic data, only chlorpyrifos, the most toxic compound (Fig. 2b), caused broad changes in protein levels. However, all but 1,2-dibromoethane altered the protein phosphorylation status in the treated larvae (Fig. 2f)— including those that are not lethal at the employed concentration (Fig. 2b). Most changes were detected in the phosphorylation pattern of proteins associated with muscle physiology, which might reflect important aspects of the stress response to xenobiotics (Extended Data Fig. 3c)^20^. In summary, the experiments described here suggest that sublethal doses of these chemicals can have pervasive effects on larval populations.

**Fig.2:**
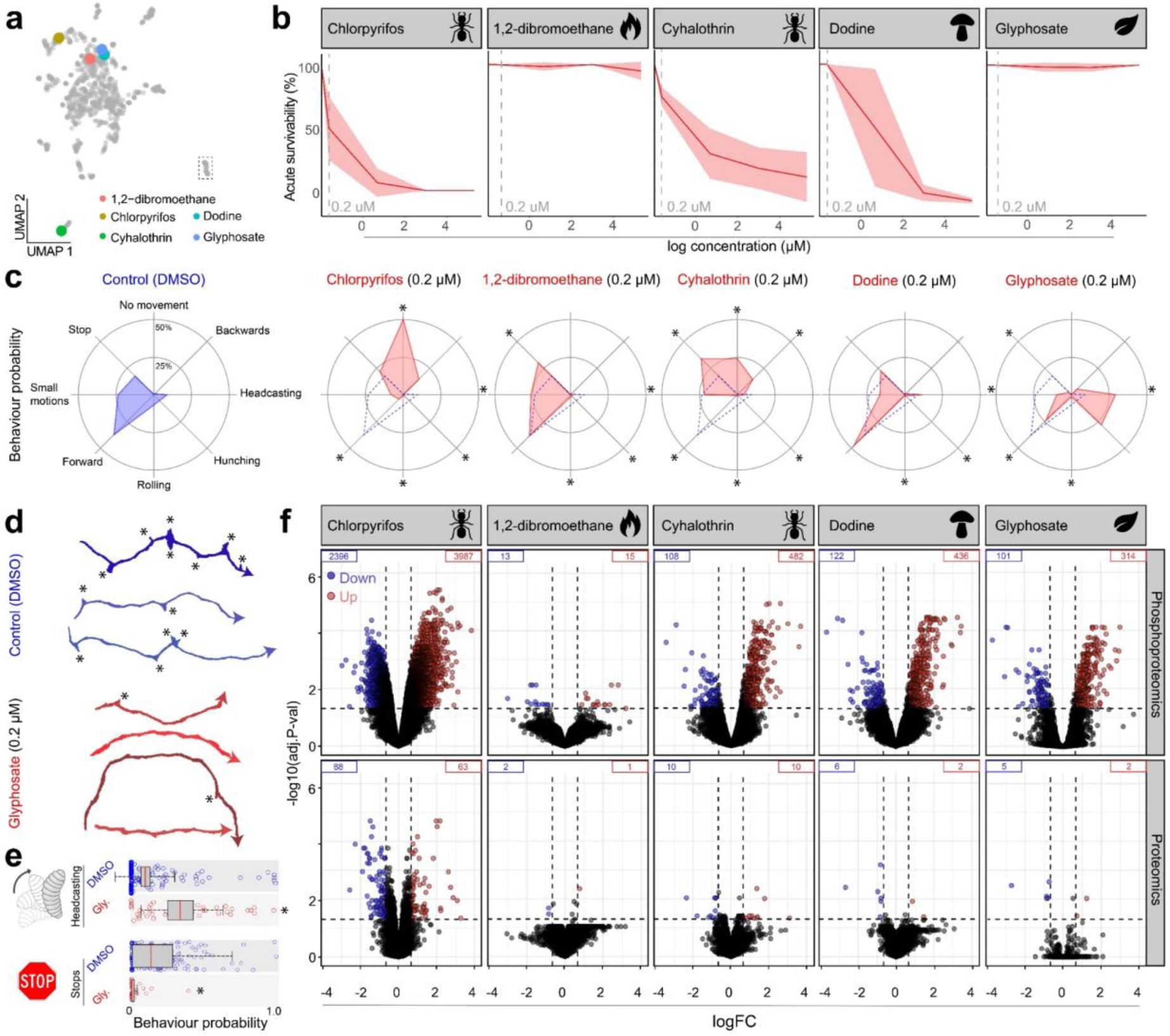
Different types of pesticides induce pervasive changes in the phosphoproteome of treated larvae. **a,** Pesticides that were selected for phosphoproteomic analysis. Their position in the UMAP described in Fig 1e is highlighted in colors. **b,** Survivability dose-response curves for these five compounds. The concentration used for the phosphoproteomic characterization of the treated larvae—0.2 μM—is highlighted by the gray dashed line. N=2, n∼25. **c,** Radar charts describing the behavioral repertoire of control (blue) or treated (red) larval populations. Each axis represents the frequency of stereotypic behaviors, noting significant changes, * p<0.05, Tukey’s test, one-way ANOVA. N=2, n∼25. **d,** Examples of the trajectories of control (blue) or glyphosate-treated (red) larval populations. Each line depicts the path of a different larva. * points out the stops. **e**, Frequencies of headcasting and stops in control (blue) or treated (red) populations. Each dot represents a different larva. * p<0.05, Tukey’s test after one-way ANOVA. N=2, n∼25. **f,** Volcano plot analysis showing significant changes in the phosphoproteome (upper row) and proteome (bottom row) of larvae treated overnight with different pesticides (column headers) at 0.2 μM. Y-axis represents the significance of the changes as minus log transformed adjusted P value. Horizontal dashed lines mark the p<0.05 threshold. X-axis indicates the magnitude of the changes as log-transformed fold change. Vertical dashed lines mark the FC>1.5 threshold. The number of annotations for which significant changes were detected is shown within each panel (down in blue, up in red).

### Higher temperatures amplify the effects of agrochemicals

The high-throughput platform described here enabled us to study not only the physiological effects of sublethal concentrations of pesticides, but also how pesticides can interact with other factors that may be driving the decline of insect populations, such as global warming^1,2^. For these experiments, we selected a subset of 49 pesticides (Table S2) showing high environmental prevalence in Europe (KEMI database)^21^ or in the USA (data from the EPA^22^ and the US geological survey^23^). We purposely chose these chemicals with the aim of being inclusive across chemical identities and pesticide types encompassed in the main library (Fig. 3a).

**Fig. 3:**
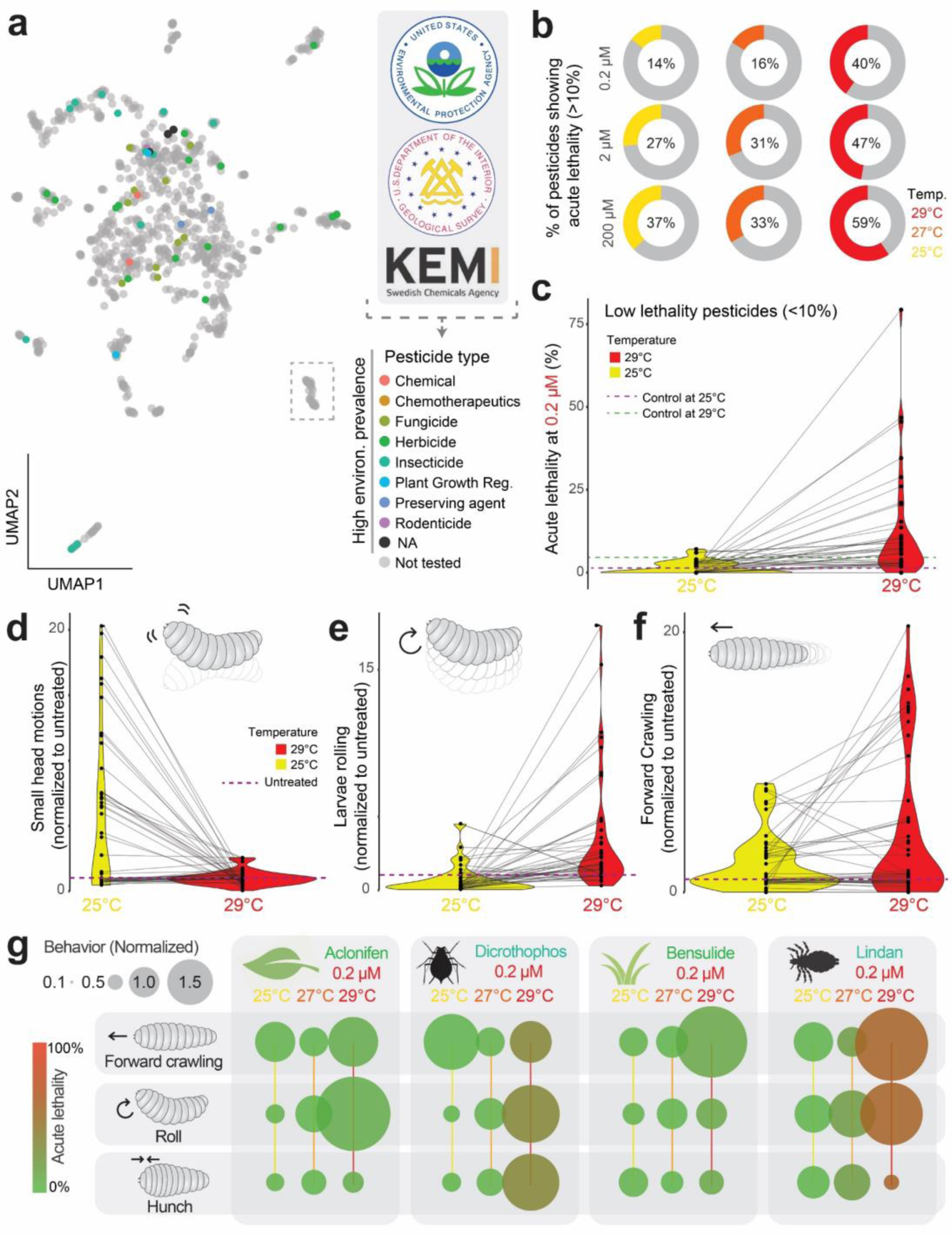
Higher temperature increases acute lethality and modulates agrochemical-induced behaviors. **a,** Subset of agrochemicals selected based on their high environmental prevalence (according to data from KEMI, EPA, and the US geological survey) for their use in subsequent experiments. These molecules are color-coded based on their agrochemical type on top of the UMAP described in Fig 1e. **b,** Percentage of agrochemicals that lead to acute lethality (>10% in 2 replicates) at three different concentrations and three different temperatures. **c**, Average of acute lethality (N=2, n∼20) for pesticides considered non-lethal (<10%) at 25^◦^C. Each dot represents the average of a different molecule. Lines connect the values observed for the same molecula at different temperatures. **d-f**, Average frequencies (N=2, n∼20) of different stereotypic movements (small head motions in **d**, rolling in **e**, and forward crawling in **f**) in populations treated with the pesticides highlighted in **a**. The frequencies are normalized to measurements from control populations at the indicated temperature. **g**, Examples of temperature-dependent behavior alterations induced by agrochemicals. The radius of each circle is proportional to the average normalized frequency (N=2, n∼20) of the indicated behavior. The color code shows the average acute lethality (N=2, n∼20) measured for each condition.

We next exposed larval populations to each of these 49 pesticides employing the same pipeline but increasing the overnight incubation temperature. Even though higher temperatures did not significantly affect larval survivability after a 16-hour exposure (Extended Data Fig 3a), we found that about two times more compounds induced acute lethality when the experiments were conducted at 29^◦^C compared to 25 or 27^◦^C (Fig. 3b and Extended Data Fig. 3b-l). Hence, many pesticides that showed low lethality (<10%) at 25^◦^C started exhibiting significantly higher lethality when the environmental temperature was increased by just four degrees (Fig. 3c, p=0.300 at 25^◦^C vs p=0.008 at 29^◦^C, two-tailed T-test). For example, the insecticide lindan, which was not lethal when used at 0.2 μM at 25^◦^C (0% acute lethality), became strongly lethal (79% acute lethality) at 29^◦^C. Additionally, while many of these molecules did change the behaviors of the treated populations when administered at 25^◦^C, the alterations triggered at 29^◦^C were, in many cases, radically different (Fig. 3d-f and Extended Data Fig. 3m-p). It should be noted that some of these molecules, such as the herbicides aclonifen and bensulfide, dramatically altered behavior without impacting acute lethality (Fig 3g). Such behavioral effects would likely go undetected when employing tests that rely exclusively on lethality^6^.

### Combinations of agrochemicals can affect life-history traits at sublethal concentrations

Pesticides are often used in combinations for industrial agriculture^24^. Additionally, the environmental persistence of certain molecules can lead to animals being exposed to two or more pesticides simultaneously^25,26^, even if the chemicals were not designed for concurrent use. Previous studies have reported non-additive effects of agrochemicals on arthropod systems such as *Daphnia*^27^ and honeybees^28^, but how frequently these combinatorial effects manifest across different classes of chemicals remains unclear.

To test possible combinatorial effects of the chemicals, we exposed larval populations overnight to all possible pair-wise combinations of 22 highly prevalent pesticides (Fig. 3a) at the lowest concentration used in the previous assay, 0.2 μM, and measured acute and long-term lethality, as well as the behavioral states of the populations. We find evidence for non-linear interactions between agrochemicals across many combinations (Extended Data Fig. 4a-b), suggesting synergistic pesticides interactions may be a widespread phenomenon. Synergy on acute lethality was confirmed for two of these combinations through a checkerboard analysis (Extended Data Fig. 4c-d).

Field studies often detect more than a couple of agrochemicals in the sampled sites^29^. Therefore, to test the sublethal effects of pesticides in more realistic environments, we focused on a single pesticide mix composed by nine pesticides (Table S3) that were detected in air samples across Germany^29^ (Fig. 4a-b). Following an overnight exposure to these combined molecules at 0.02 μM—well within standard environmental prevalence ranges of these chemicals^13,30^—we observed neither acute lethality (Fig. 4b) nor behavioral alterations (Extended Data Fig. 5a). However, when the larvae were fed solid food containing this pesticide mix from hatching up to the 3^rd^ instar, we found widespread changes in behaviors (Fig. 4c, Extended Data Fig. 5b). We found that these treated populations were developmentally delayed, requiring approximately one additional day to pupate (*p*<0.001, two-tailed T-test, Fig. 4e). Moreover, the adults emerging from these pupae showed a significantly reduced egg laying rate (*p*<0.001, two-tailed T-test, Fig. 4f). Remarkably, none of the agrochemicals included in the mix are labeled as insecticides (Table S3). Taken together, these findings demonstrate that—even under sublethal conditions and at concentrations found in natural habitats—pesticides can compromise the fitness of insect populations.

**Fig. 4:**
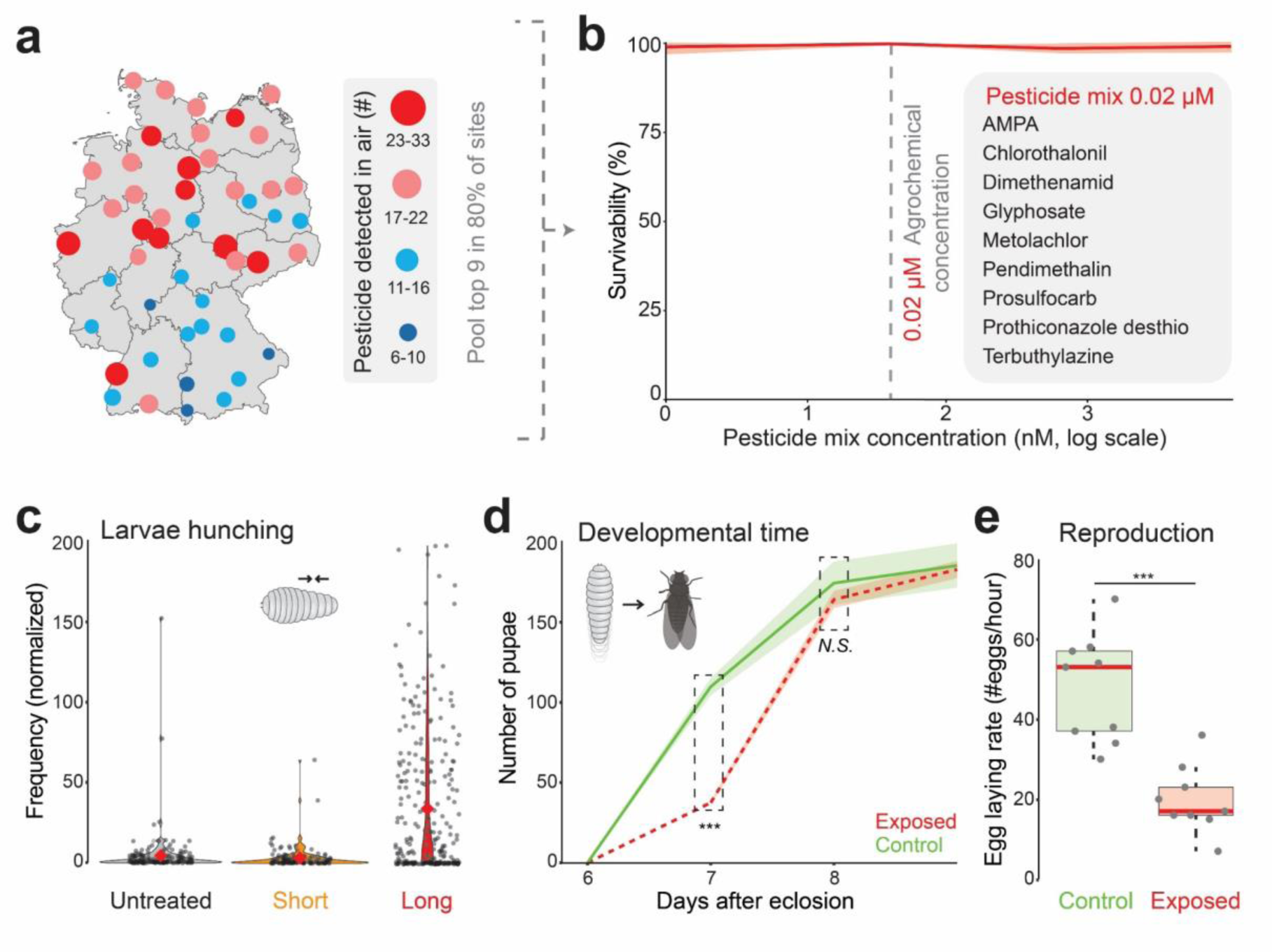
Long-term exposure to pesticide mix reveals changes in life-history traits. **a**, Map of Germany showing the number of pesticides detected in the air (based on Kruse-Plaß et al.^29^) The number of detected pesticides in each site is depicted in the color code and the radius of each circle. **b**, Average (N=2, n∼20) survivability (percentage of surviving larvae) after overnight exposure to the noted pesticide mix at different concentrations. **c**, Hunching frequencies (normalized) of larval populations exposed to the 0.02 μM pesticide mix for 16 h (short) or for 5 days (long). Red dots show the mean across all molecules. **d**, Accumulated number of pupae observed each day after hatching, on populations (N=3) of 200 larvae exposed to the pesticide mix at 0.02 μM from eggs onwards. ***p<0.01, two-tailed T-test. **e**, Number of eggs laid per hour in populations (N=9) of 100 flies (50 females) treated with the pesticide mix at 0.02 μM from eggs onwards. Red dots show the mean across all molecules. ***p<0.001, Wilcoxon test. The center line is the median, and the boxed region represents the interquartile range.

### Natural isolates show lower sensitivity, specifically to organophosphates

To control for inbreeding depression in our lab strains^31^, we next tested two other *D. melanogaster* lab strains (CantonS and Ind), as well as two different fly species, *D. willistoni* and *D. virilis* (Extended Data Fig. 7a-d). In addition, to probe if populations may adapt to environmental agrotoxins we established two iso-female lines from *D. melanogaster* flies collected near Naples, Italy—a site with detectable levels of organophosphate pesticides^32^. We compared the sensitivity of these strains with our standard lab stock by exposing them to the 49 highly prevalent pesticides described above (Fig. 3a and Table S2). While resistance profiles across most classes of chemicals were similar among the different tested lines, we found that both natural isolates showed greater resistance to organophosphates than our lab strains (Extended Data Fig. 7a-d). These results highlight that it is possible to identify subtle differences in sensitivity among insect populations, possibly due to adaptive responses to chemical exposure.

### Behavior is also affected by sublethal agrochemical doses in mosquitoes and butterflies

Next, we explored if our approach could be used to detect the sublethal effects of pesticides on the behavior of other insects. We focused on medically- and economically-relevant species, such as a malaria vector, the mosquito *Anopheles stephensi,* and a widely distributed pollinator, the butterfly *Vanessa cardui* (Fig. 5). In the case of mosquitoes, we exposed larval populations overnight to varying concentrations of a neonicotinoid (thiacloprid), a pyrethroid (cyhalothrin) and a fungicide (dodine) on multiwell plates, followed by recording and tracking of these larvae in order to quantify their movement patterns (Fig. 5a-d). Notably, for all pesticides, we found larvae were moving significantly slower at concentrations associated with negligible or low lethality (Fig. 5e-p). In the case of *V. cardui*, we exposed 4^th^ instar caterpillars to solid food containing these same three molecules. The movement of the caterpillars was then recorded in agar plates, and their paths were again tracked (Fig. 5q-s). Interestingly, while only cyhalothrin showed some level of lethality within the range of concentrations employed in this assay (Fig. 5t), all three molecules affected the movement patterns of the treated caterpillars (Fig. 5u-f’). Thus, these results highlight that sublethal concentrations of pesticides can also affect the behavior of species with high ecological, economic, and clinical relevance.

**Fig. 5:**
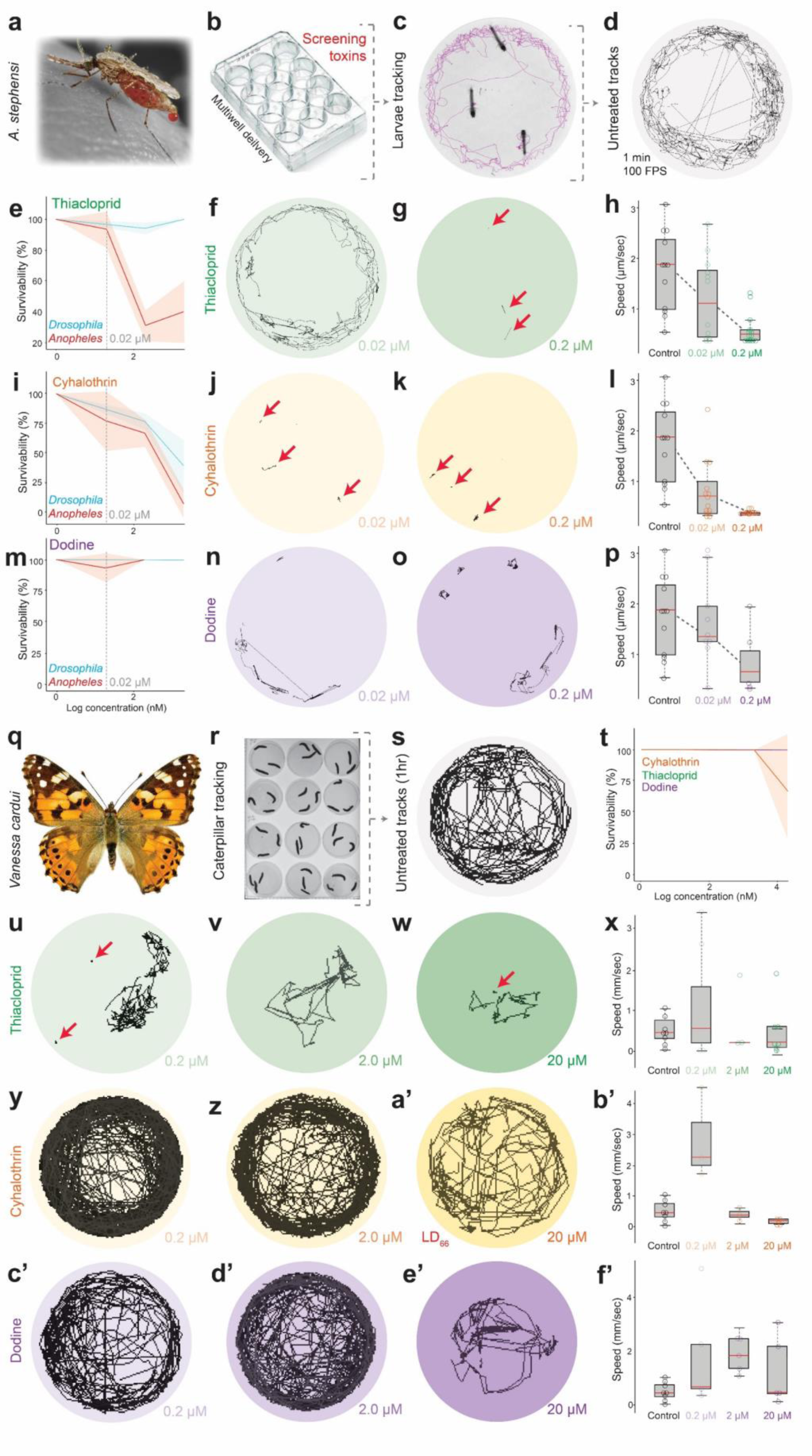
Sublethal effects of pesticides on the behavior of mosquitoes and butterflies. **a-d**, Schematic of the experimental design used for analyzing the impact of pesticides on the behavior of mosquito larvae. Fourth instar larvae of *Anopheles stephensi* (**a**) were exposed overnight to different pesticides at either 0.2 or 0.02 μM in multiwell plates (**b**). The larvae were then recorded in each well and tracked using Mtrack2 (**c, d**). **e,i** & **m**, Survivability curves of *Drosophila* and *Anopheles* larvae exposed overnight to Thiacloprid (**e**), Cyhalothrin (**i**), or dodine (**m**) in a liquid environment. **f-h**, Effects of Thiacloprid on the trajectories (**f, g**) and speed (**h**) of *Anopheles* larvae. **j-l**, Effects of Cyhalothrin on the trajectories (**j, k**) and speed (**l**) of *Anopheles* larvae. **n-p**, Effects of dodine on the trajectories (**n, o**) and speed (**p**) of *Anopheles* larvae. N=3, n=3. **p-s**, Schematic representation of the experimental design used for analyzing the effect of pesticides on the behavior of caterpillars. Fifth instar *Vanessa cardui* caterpillars (**q**) were exposed overnight to different pesticides at 20, 0.2 or 0.02 μM in solid food. The larvae were then transferred to agar plates, recorded (**r**) and then tracked using Mtrack2 (**s**). **t**, Survivability curves of *Vanessa* caterpillars exposed overnight to Thiacloprid (red), Cyhalothrin (green) or dodine (purple) in a liquid environment. **u-x**, Effects of Thiacloprid on the trajectories (**u-w**) and speed (**x**) of *Vanessa* caterpillars. **y-b’**, Effects of Cyhalothrin on the trajectories (**y-a’**) and speed (**b’**) of *Vanessa* caterpillars. **c’-f’**, Effects of dodine on the trajectories (**c’-e’**) and speed (**f’**) of *Vanessa* caterpillars. N=3, n=3. Arrows across all panels highlight larvae that are alive but barely moving. In all box plots the center line is the median, and the boxed region represents the interquartile range.

## Discussion

For effective conservation efforts to mitigate the global decline in insect populations^1,2^, the relative effects of each potential driver must be carefully disentangled. Experimental work, where variables can be tightly controlled, offers the means to do so^7^. This becomes more important when considering the synergic relationships between chemicals^24,33,34^. Together, by leveraging tools and methods from basic science, experimental research can furnish policymakers and chemical companies with the necessary information to implement rational sustainability measurements^35^.

Here, building upon our previous attempts at deep phenotyping^36^, we developed a high-throughput phenotyping platform, which enabled us to characterize the behavioral effects of more than 1,000 agrochemicals when employed at varying concentrations. Strikingly, we found that 57% of the agrochemicals in our library exert significant effects on the behavior of *Drosophila* larvae when exposed for a short time period at sublethal concentrations (Fig. 1k). While further research is needed to investigate the longer-term impact on the population fitness, our experiments revealed that key traits—such as egg laying rates—are significantly reduced by the exposure to some of these molecules at concentrations orders of magnitude under sublethal concentrations (Fig. 4). Additionally, higher temperatures increased pesticide-induced lethality and behavioral alterations (Fig. 2), highlighting the need for chemical testing under more realistic environmental conditions, especially given rising global temperatures. Together, these observations emphasize the potential for ecological developmental biology towards discussions on sustainability, as it can provide quantitative information across life stages on how biological systems react to human-driven environmental perturbations^35^.

Our findings highlight that many agrochemicals with high environmental prevalence can induce behavioral changes across insect species, even at sublethal levels. For example, dodine, a guanidine fungicide with no reported mechanism of action, which is currently sprayed in the US at 20 mM^37^, induced broad changes in the phosphoproteome of *Drosophila* larvae. Consistent with these changes, dodine altered larval behaviors in flies, mosquitoes and butterflies (Fig 2c and Fig 5) when used at concentrations several orders of magnitude below the spraying concentration (EPA report^22^). Similarly, glyphosate led to strong behavioral alterations and changes in the larval phosphoproteome when used at sublethal concentrations. In order to mitigate the environmental impact of such molecules, our findings suggest that the next generation of pesticides should be subjected to more comprehensive testing focused on sublethal effects across different representative species. Importantly, these types of assays provide more precise data on how to target pest control for medically important vector species without negatively affecting overall insect biodiversity.

The reduced egg laying rate detected in fly populations exposed to pesticides reveals that even under sublethal conditions and at concentrations found in natural habitats, these molecules can compromise the fitness of insect populations. As such, we provide empirical evidence supporting the role of agrochemicals as a driver of the collapse of insect populations, which may be further exacerbated by climate change^2,33,38^. We anticipate that assays on chemical safety inclusive of fitness parameters outside of lethality will contribute to improving chemical safety assessment to better protect the environment, secure food supplies, and safeguard animal and human health.

## Methods

### Fly strains

Unless specified otherwise, all experiments were carried out in a *w1118*;; genetic background. Natural isolates were captured in Naples (STR1) and Ischia (BTI1-2) in the summer of 2019, following a standard protocol^39^. The isofemale lines were derived from single females and had been kept in the lab for 3 years (2019-2022) before the experiments. The flies were verified as *D. melanogaster* without *Wolbachia* infection through PCR with primers described in Faria and Sucena (2017)^40^.

All fly strains were kept at standard laboratory conditions at 25^◦^C.

### Chemical library

A chemical library of 1024 agrochemicals was assembled by purchasing each molecule individually (Sigma Aldrich) and solubilizing in DMSO at a uniform concentration of 10 mM. The library was selected to encompass a comprehensive range of pesticide types (eg: herbicides, insecticides, fungicides etc), and also to represent a wide variety of usages (eg: active-use pesticides, obsolete and banned pesticides, pesticide precursor chemicals, pesticide transformation products). Full details are given in Table S1.

### Molecular fingerprint analysis

Molecular fingerprints were generated from chemical compounds represented in the Simplified Molecular Input Line Entry System (SMILES) format using the rdkit.Chem.MolFromSmiles function in RDKit Python package (version 2022.09.1). Two types of molecular fingerprints were calculated based on the SMILES representations of the molecules in the chemical library, namely Morgan and RDKfingerprint using AllChem.GetMorganFingerprintAsBitVect and rdkit.Chem.RDKFingerprint functions from RDKit, respectively. Following this, Uniform Manifold Approximation and Projection (UMAP) algorithm^15^ was utilized from the umap-learn python package (version 0.5.3) to reduce the dimensionality of the chemical space for both types of fingerprints. Three clustering algorithms from scikit-learn python package (version 1.1.3) —k-means, bisecting k-means, and agglomerative clustering—were evaluated, with varying numbers of clusters explored for each algorithm. K-means clustering of UMAP dimensionally reduced Morgan fingerprints yielded the lowest Davies-Bouldin score with compounds separated into 11 clusters.

### Pesticide delivery and video recording

Pesticide delivery was carried out according to the protocol described by Gasque et al^9^. Briefly, five days old larvae (3^rd^ instar) were transferred to 24-well multiwell plates containing in each well a different agrochemical at the desired concentration in a liquid medium (yeast extract 10% m/v, glucose 10% m/v and sucrose 7.5% m/v, without antimycotics or antibiotics). For the controls, only the solvent (DMSO) was added to the well. Between 15 and 30 larvae were added to each well. Larval density was controlled in the original populations to minimize metabolic heterogeneities at the onset of the experiments.

The multiwell plates were sealed with a breathable membrane and kept at 25^◦^C (unless otherwise noted, see Fig. 3) for 16 h, before counting the fraction of surviving larvae. Then the larvae were transferred to the agar plates, where they were recorded for 1 min at 30 fps using an FL3-U3-13Y3M-C CMOS camera. Finally, the larvae were transferred to vials with standard food (solid, pesticide-free), and the number of adult flies was counted 10 days later. Adults that did not emerge from the puparium after this period (15 days after egg laying) were considered dead.

The original videos together with the processed files used for the analysis are publicly available at the BioImage Archive (accession number S-BIAD970).

### Image analysis and behavior characterization

In one pipeline (Larva-Tagger), the videos were first processed using Multi-Worm Tracker^41^, a package that extracts low-level features including the position and outline of each larva. More specifically, the mwt-core C++ online tracking library was combined with the Choreography Java post-processing utility to track the larvae. Due to the high number of video files to process, the selection of tracking hyper-parameters (pixel and size thresholds) was automated using the Optuna Python optimization library and targeting a number of tracked objects as close as possible to the known number of larvae in the assay. The resulting moving objects, as identified by the tracking procedure, were further post-processed on a per-assay basis, rejecting the moving objects whose average surface area was more than twice the median area across all moving objects, or less than half the median area. Additionally, to crop the tracks that collapsed to a small blob prior to disappearing, a change-point detection algorithm was applied to determine from what time step to reject the end part of the track.

As a second processing step after the tracking procedure, the resulting parameters were used to explore larval behavior using LarvaTagger, a piece of software that relies on a deep neural network for stereotypical behavior identification., The stereotypical actions or postures included: forward and backward crawl, roll, bend/head cast, hunch, stop, as well as a “small action” category of unidentified actions or postures.

For statistical testing, any registration that included a fraction of dead larvae greater than 10% of the entire dataset was removed. Additionally, artefactual signal was avoided by removing any lineage (single larva) that was tracked for fewer than 10 frames.

All the code is free and open source. The full tracking and behavior tagging pipeline is available at https://gitlab.com/larvataggerpipelines/Pesticides. For the most part, this code is specific to the computer cluster at EMBL Heidelberg. Its more-generally applicable components include mwt-container, available at https://gitlab.com/larvataggerpipelines/mwt-container, for the head-less tracking of larvae in avi video files. This package provides a Docker image (flaur/mwt-container:0.1 pulled from quay.io) that itself ships with mwt-core (also available at https://github.com/Ichoran/mwt-core), Choreography (also available at https://github.com/Ichoran/choreography) and the mentioned optimization procedure (also available as a separate package at https://github.com/Lilly-May/larva-tagger-tune). The above-mentioned post-processing steps are implemented in Julia and are part of the Pesticides repository. The behavior tagging software is available at https://gitlab.pasteur.fr/nyx/LarvaTagger.jl. Docker images are also available. For the present analysis, the flaur/larvatagger:0.16.4-20230311 image pulled from hub.docker.com was used.

In the second pipeline (Trajectories), data was preprocessed to reduce the frame rate by a factor of 15 which is sufficient for the tracking task, then the value range was inverted and well boundaries were masked with an automatic algorithm. Larvae were then segmented using random forest-based Pixel Classification in ilastik^42^ on 36 randomly selected images. The random forest classifier was trained interactively with 613 and 1662 annotated pixels for foreground, and background, respectively, using all available default features. The resulting probability maps were thresholded at 0.5 for use in Tracking^43^ in ilastik. Division and Object-Count classifiers were trained interactively using default tracking features. We augmented the training data for tracking by adding time frames 0, 5, 10, and 15 from 39 datasets. This was done in order to improve count estimates for under-segmented agglomerations of larvae, often present at the beginning of the time series. An R script was then used to compute 9 trajectory features from the sets of larvae coordinates. These trajectory features are displacement (distance between first and last larva positions), distance travelled (sum of the distances travelled between consecutive frames), straightness (displacement over distance travelled), convex hull perimeter, convex hull area, wandering (distance travelled over convex hull perimeter), curvature (sum of the angles between directions on consecutive frames), radius of gyration, aspect.ratio (ratio of the axes of the ellipse fitted to the larva positions). Features were then aggregated by taking the median values for each well. The preprocessing, pixel classification and tracking tasks are implemented as nextflow workflows and code is available in the project’s repository: https://git.embl.de/grp-crocker/agrotoxin.

The trajectories shown in Fig 2d were generated using FIMtrack^44^.

### Statistical analysis and hits identification

The LarvaTagger tag sequences were first transformed into frequency tables, a probability vector was computed by counting the occurrence of each tag and dividing them by the total number of tags. One pseudocount was added to each tag to avoid having probability equal to 0 for any tag, so that logarithmic operations could be performed on the values in the subsequent step. Behavioural frequencies were converted into information content values using the Shannon *Information Content* formula, where *f* is the frequency of a tag in the probability vector, and *p* is the median probability of a tag in the control population:

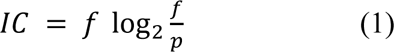

While values of a probability vector are compositional and thus only convey relative information, IC values are independent from one another and can be used for standard statistical analysis. For subsequent analysis, the values were aggregated by taking averages for each well.

For hit detection, we expect that conditions affecting larval behavior would produce features distributions that differ from the distribution in the control population and thus can be detected as statistical outliers. A classical multivariate outlier detection method makes use of the Mahalanobis distance such that objects with a high Mahalanobis distance to the control mean are considered outliers. The Mahalanobis distance of each well from the distribution observed in the control population was computed. Since the squared Mahalanobis distance follows a Chi-squared distribution, we used this property to associate a p-value to each condition. We consider a condition as affecting larval behavior if its FDR-adjusted p-value is below 0.05..

Positive hits of the screen were then defined as those conditions that significantly affected larval behavior in at least the median number of replicates, for most conditions, this corresponds to a minimum of 2 out of 3 replicates.

Trajectory features and Larva-Tagger classes were processed separately yielding two sets of partially overlapping hits (intersection over union of 35%)

The code used for this analysis is available in the project’s code repository: https://git.embl.de/grp-crocker/agrotoxin.

### Phosphoproteomics

For the (phospho)proteomics experiments, the lysis buffer was composed of 4 M guanidinium isothiocyanate, 50 mM 2-[4-(2-hydroxyethyl)piperazin-1-yl]ethanesulfonic acid (HEPES), 10 mM tris(2-carboxyethyl)phosphine (TCEP), 1% N-lauroylsarcosine, 5% isoamyl alcohol and 40% acetonitrile adjusted to pH 8.5 using 10 M NaOH. For sample lysis, 500 μL of lysis buffer was added to 50 larvae before performing bead beating with 1.6 mm zirconium silicate beads for 30 seconds at 6 m/s (Bead ruptor elite - Omni international). Samples were then centrifuged at 16,000 × g at room temperature for 10 minutes to remove cell debris and nucleic acid aggregates. Protein concentrations were determined using tryptophan fluorescence assay as described in Wisńiewski et al.^45^. Then, ice-cold acetonitrile was added to the samples to induce protein precipitation to a final concentration of 80% acetonitrile. After 10 minutes, the samples were centrifuged to remove the solution, and protein precipitates were washed two times with 200 µL 80% acetonitrile and two times with 200 µL 70% ethanol (1,000 × g for 2 minutes for each step). Finally, the digestion buffer composed of 100 mM HEPES pH 8.5, 5 mM TCEP, 20 mM chloroacetamide and trypsin (TPCK treated trypsin, Thermo Fisher Scientific) was added to the protein precipitates. The ratio of trypsin:protein was fixed to 1:25 w/w and the maximum final protein concentration to 10 µg/µL. Tryptic digestion was carried on overnight at room temperature under mild shaking (600 rpm). After digestion, samples were acidified to 1% TFA and desalted using Sep-pak tC18 columns (Waters), eluted using 0.1% TFA in 40% acetonitrile, and dried using a vacuum concentrator. Aliquots corresponding to 5% of the peptides were saved for the full proteome measurements before phosphopeptides enrichment.

Prior to enrichment, lyophilized peptides were resuspended in buffer A (80% ACN, 0.07% TFA), sonicated and centrifuged at 16,000 × g. The enrichment was performed as described in Leutert et al.^46^ using the KingFisher Apex robot (Thermo Fisher Scientific) and 50 μL of Fe-NTA Magbeads (PureCube) per sample. After 5 washes with buffer A, bound phosphopeptides were eluted by the addition of 100 µL of 0.2% diethylamine in 50% acetonitrile before lyophilization.

For TMT labeling, 15 µg of dried non-modified peptides and the totality of enriched phosphopeptides were resuspended in 10 μL 100 mM HEPES pH 8.5 before addition of 4 μL or 2 μL of TMT reagent at a concentration of 20 μg/μL in acetonitrile in the case of non-modified and phosphopeptides, respectively. After 1h at room temperature, the labeling reaction was quenched for 15 minutes with the addition of 5 μL of 5% hydroxylamine. Labeled peptides belonging to the same experiment were then pooled and subsequently lyophilized. Prior to fractionation, the full proteome samples were desalted using Sep-pak tC18 columns (Waters), while the labeled phosphopeptides were resuspended into 50 µL of 10% TFA and desalted using C18 stagetips^47^ made in-house and packed with 1 mg of C18 bulk material (ReproSil-Pur 120 C18-AQ 5 µm, Dr. Maisch) on top of the C18 resin plug (AttractSPE disks bio - C18, Affinisep).

Off-line fractionation was performed on an Ultimate 3000 Liquid Chromatography system (Thermo Fisher Scientific). Samples were reconstituted in 18 μL buffer A (0.05% TFA in MS grade water with 2% acetonitrile), and the peptides were separated on a Hypercarb column (100mm, 1.0mm ID, 3µm particle size, Thermo Fisher Scientific) at a temperature of 50°C and a flow rate of 75 µL/min. The linear separation gradient started 1 minute after injection and increased from 13% buffer B (0.05% TFA in acetonitrile) to 42% buffer B after 95 minutes, before increasing to 80% buffer B in 5 minutes. The column was washed with 80% buffer B for 5 minutes before being re-equilibrated for 5 minutes with 100% buffer A. Fractions were collected from 4.5 minutes to 100.5 minutes with a 2 minutes collection period, resulting in 48 fractions which were concatenated into 24 fractions (n fraction being pooled with the n + 24 fraction). Samples were dried using a vacuum concentrator prior to LC-MS/MS analysis.

Peptides were separated using an UltiMate 3000 RSLCnano system (Thermo Fisher Scientific). Peptides were first trapped on a cartridge (Precolumn; C18 PepMap 100, 5 μm, 300-μm i.d. × 5 mm, 100 Å) before separation on an analytical column (Waters nanoEase HSS C18 T3, 75 μm × 25 cm, 1.8 μm, 100 Å). Solvent A was 0.1% formic acid in LC–MS-grade water and solvent B was 0.1% formic acid in LC–MS-grade acetonitrile. Peptides were resuspended in a loading buffer constituted of 50 mM citric acid, 1% trifluoroacetic acid and 2% acetonitrile before loading onto the trapping cartridge (30 μL/min solvent A for 3 min) and eluted with a constant flow of 300 nL/min using an analysis time of 120 minutes (full proteome samples: linear gradient of 10–28% B for 99 minutes ; phosphopeptide samples: linear gradient of 9-26% B for 99 minutes). The linear gradient was followed by an increase to 40% B within 5 minutes before washing at 85% B for 4 minutes and re-equilibration to initial conditions.

The LC system was coupled to an Exploris 480 mass spectrometer (Thermo Fisher Scientific) operated in positive ion mode with a spray voltage of 2.2 kV and a capillary temperature of 275 °C. The mass spectrometer was operated in data-dependent acquisition (DDA) mode with a maximum duty cycle time of 3 seconds. Full-scan MS spectra with a mass range of 375– 1,500 m/z were acquired in the Orbitrap using a resolution of 90,000 with a maximum injection time of 45 ms and automatic gain control (AGC) set to 3 × 106 charges. The most intense precursors with charge states 2–7 and a minimum intensity of 2 × 105 were selected for subsequent HCD fragmentation (isolation window of 0.7 m/z and normalized collision energy of 32%) with a dynamic exclusion window of 30 seconds or 25 seconds for the full proteome and phosphopeptide samples, respectively. MS/MS spectra were acquired in profile mode with a resolution of 45,000 in the Orbitrap (maximum injection time of 100 ms, AGC target of 2 × 105 charges and first mass fixed at 110 m/z).

### Phosphoproteomics – data analysis

Mass spectrometry raw files were converted to mzmL format using the MSConvert software from Proteowizard^48^ using peak picking from the vendor algorithm and keeping the 300 most intense peaks. Files were then searched using MSFragger v3.8 in Fragpipe v19.1^49^ against the *Drosophila melanogaster* proteome (with one protein per gene, downloaded from Uniprot) including known contaminants and the reversed protein sequences. The default TMT16 and TMT16-phospho MSFragger workflows were used and search parameters were optimized during the first search. Modifications were set as follow: fixed modifications = carbamidomethyl on cysteine and TMTPro on lysine; variable modifications = acetylation on protein N termini, oxidation of methionine and TMTPro on peptide N termini, as well as variable phosphorylation of serine, threonine and tyrosine residues with a maximum of 3 phosphorylation events per peptide. Phosphorylation sites localization probabilities were calculated using PTMProphet^50^ and the TMT reporter ion intensities were determined by TMT-Integrator. The psm.tsv files were used for subsequent data analysis.

First, the median was computed to consolidate peptide spectrum matches into phosphopeptides. Following this, intensity matrices were filtered to eliminate proteins and phosphopeptides with zero intensity values. Subsequently, the phosphoproteomic matrices were adjusted to account for changes in protein abundance. This involved calculating a normalization factor for each phosphopeptide, expressed as the ratio of the median intensity of the phosphopeptide relative to the median intensity of its parent protein.

After this correction, both intensity matrices were normalized using the vsn R package (v3.68.0). Differential abundance analyses were then performed by comparing treated and control samples, utilizing the limma R package (v3.56.2). To categorize a protein or phosphopeptide as upregulated or downregulated, specific criteria were applied: an adjusted P value < 0.05 and an absolute fold change > 1.5.

To explore the functional significance of these findings, Gene Ontology enrichment analyses were conducted for the list of hits associated with each treatment. These analyses were carried out using the ClusterProfiler R package (v4.8.2) and the org.Dm.eg.db R package (v3.17.0). Fisher’s exact test was specifically utilized, with the background set comprising all proteins identified in at least one of the studies, encompassing both proteins and phosphoproteins.

The mass spectrometry proteomic data have been deposited in the ProteomeXchange Consortium via the PRIDE partner repository with the dataset identifier PXD046850.

### Pupariation time and egg laying

Long term exposure to the pesticide mix was achieved by rearing the larvae on solid cornmeal food containing each pesticide at 0.02 μM. 200 eggs were transferred to each vial, and then the number of pupae in the vial wall was counted every day between the 6^th^ and 9^th^ days after the onset of the experiment.

At day 10, 100 adult flies coming from these vials (i.e. exposed to the pesticide mix from hatching onwards) were transferred to egg collection cages, and the number of eggs laid per hour (between 10 and 11 am) in apple agar plates was quantified.

### Pesticide delivery to mosquitoes and caterpillars

*Anopheles stephensi* larvae were reared under standard insectary conditions, at 28°C and 80% humidity, with a 12:12 photoperiod and an hour dawn:dusk cycle. 4^th^ instar larvae were then transferred from Heidelberg University Hospital to EMBL, Heidelberg, where they were moved to 24-well plates containing the indicated pesticide at the desired concentration in water. For the control populations, only the solvent (DMSO) was added to the well. The plates were then incubated at 25°C for 16 h, and then the larvae within each well were recorded for 2 min at 100 fps using an FL3-U3-13Y3M-C CMOS camera. Individual larvae were then tracked using Mtrack2 (v2.0.1) for ImageJ.

*Vanessa cardui* caterpillars (5^th^ instar) were bought from Insect Lore UK (https://www.insectlore.co.uk/). The caterpillars were kept for 16 h on commercial food (also provided by Insect Lore UK) supplemented with each agrochemical at the desired concentration, and then they were transferred to agar plates, where they were recorded for 1 h at 1/10 fps using a regular webcam (Logitech, 1080p).

## Acknowledgments

M.Z.K, M.S., M.Z., and J.C. are supported by EMBL. L.G. and X.C.L. are supported by a fellowship from the European Molecular Biology Laboratory Interdisciplinary Postdoc Programme (EIPOD) under Marie Sk1odowska-Curie Actions cofund (grant agreement number 847543 and 664726, respectively). V.I. and J.C. are supported through state funds approved by the State Parliament of Baden-Württemberg for the Innovation Campus Health + Life Science Alliance Heidelberg Mannheim. A.L.B and V.I. are supported by the Deutsches Zentrum für Infektionsforschung (DZIF, TTU03.705), and the Deutsche Forschungsgemeinschaft (DFG, German Research Foundation) – project number 240245660-SFB 1129. We would like to thank the EMBL Chemical Core facility for help with the chemical libraries, and EMBL’s Planetary Biology Research Theme. For comments on the project and manuscript, we thank Scott Gilbert, Claire Standley, Nicolas Frankel, Arnaud Martin, and David L. Stern. We also thank Arnaud Martin for his valuable support in butterfly breeding. We would like to thank all members of the Crocker group for their feedback and support.

