## Supplementary material for "Pervasive sublethal effects of agrochemicals as contributing factors to insect decline"

**Extended Data Fig. 1. Molecular fingerprints of the agrochemical library.**

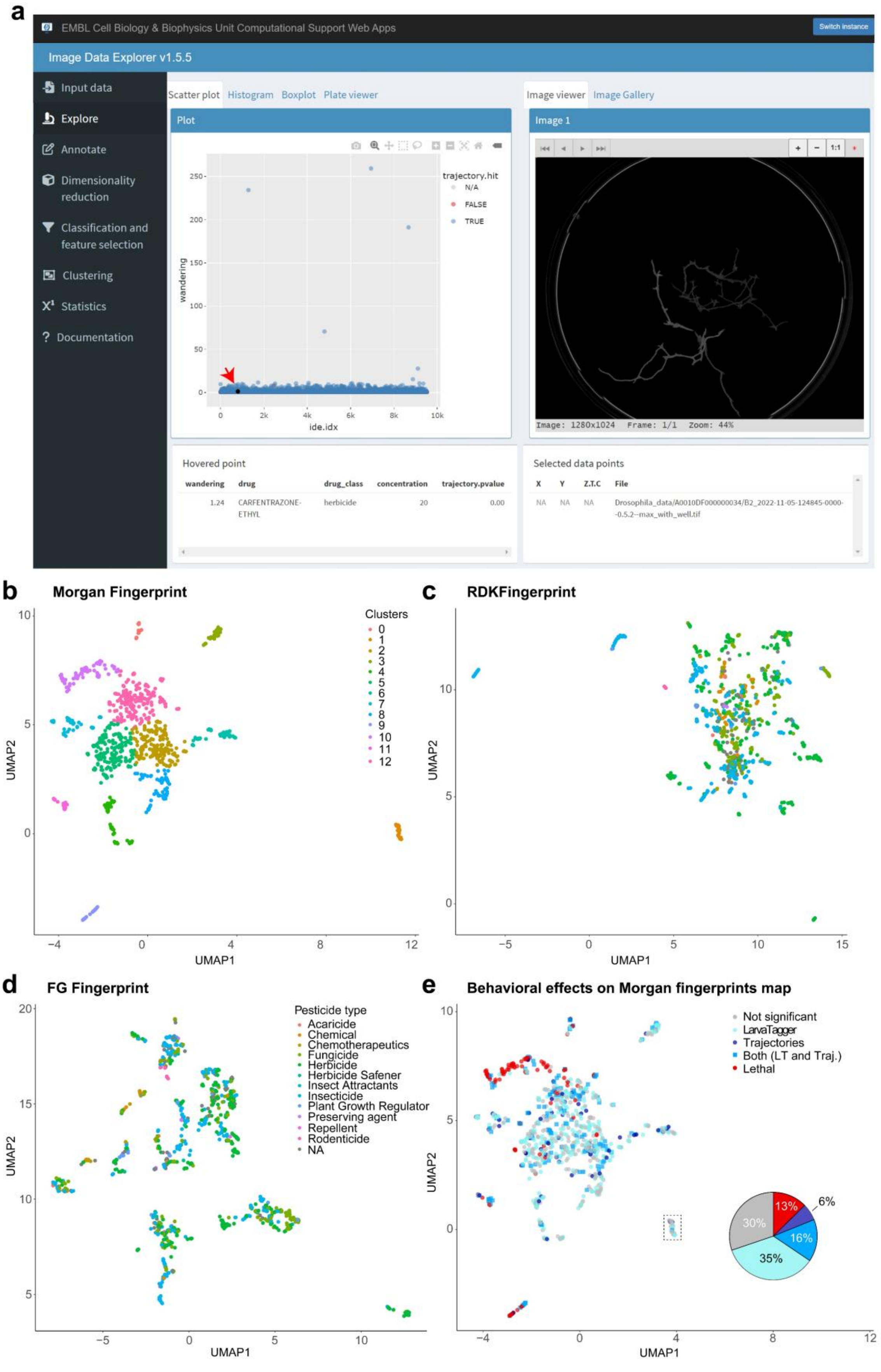

**a,** Screenshot showing the Image Data Explorer application that can be used to visualize the results of the screen through the website agrotoxin.embl.de The image on the right shows the trajectories in one replicate for the condition highlighted with the red arrow (carfentrazone-ethyl at 20 µM). **b-d**, UMAP projections of the Morgan (**b**) RDK (**c**) or Functional Groups (**d**) molecular fingerprints of the different molecules encompassed in the library. The color code shows k-means clusters (**b**) or the pesticide types (**c-d**). **e**, Classification of pesticides either as lethal at 2 µM (red), significantly ($p<0.05$ for at least the median number of replicates at that concentration) altering the frequencies of stereotypic behaviors (Larva-Tagger, light blue), trajectories (dark blue), or both (blue squares). Those with no significant effect on behavior are shown in grey. This classification is shown as a color code on top of the UMAP described in **e**.

**Extended Data Fig. 2. Acute and long-term lethality across the library.**

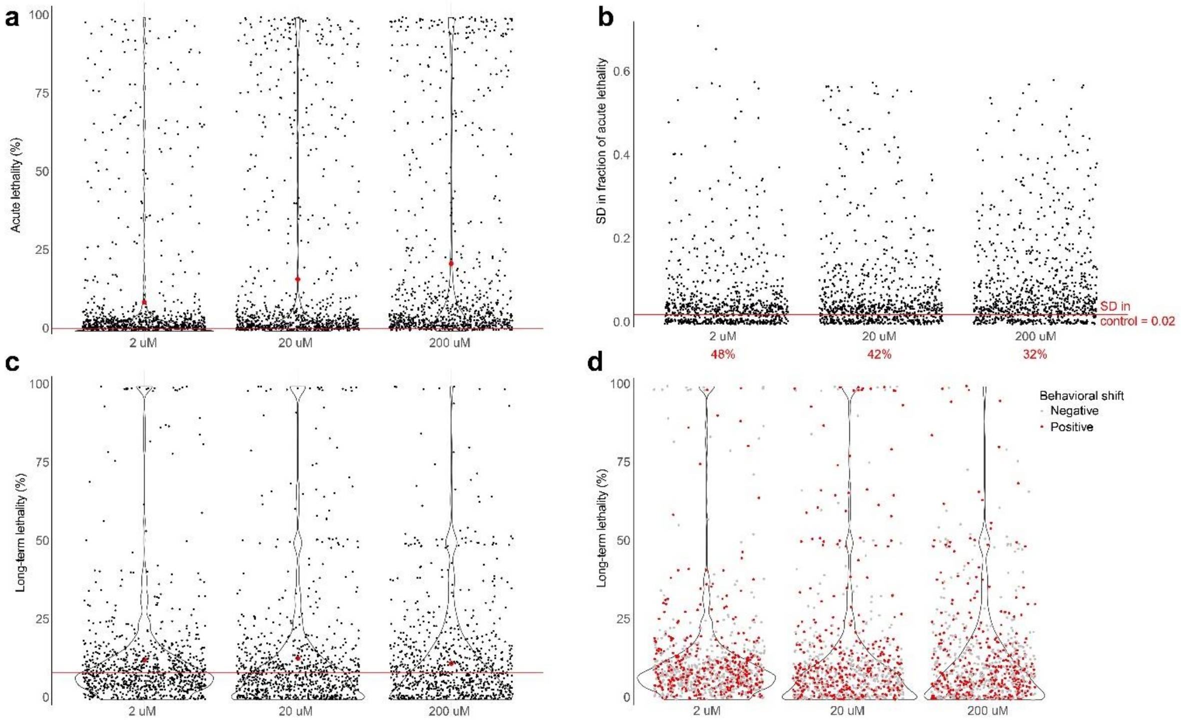


**a,** Average acute lethality (N=3, n~20) observed for all the tested pesticides at the reported concentrations. Red dots represent mean values across all molecules. The horizontal red line shows average acute lethality for control populations. **b,** Standard deviation (N=3, n~20) of acute lethality observed for all the tested pesticides when used at the reported concentration. The horizontal red line shows the SD for control populations. The red numbers at each concentration report the number of pesticides with $SD>SD_{control}$. **c,** Average long-term lethality (N=3, n~20) observed for all tested pesticides at the reported concentrations. Red dots represent mean values across all molecules. The horizontal red line shows average acute lethality for all control populations. **d,** Same plot as in c, with color-coded dots (agrochemicals) based on the effect each molecule has on larval behavior (red=significant effect on behavior; grey=non-significant).

**Extended Data Fig. 3. Gene ontology analysis of phosphoproteomic hits on larvae exposed to molecules that induce behavioral alterations.**

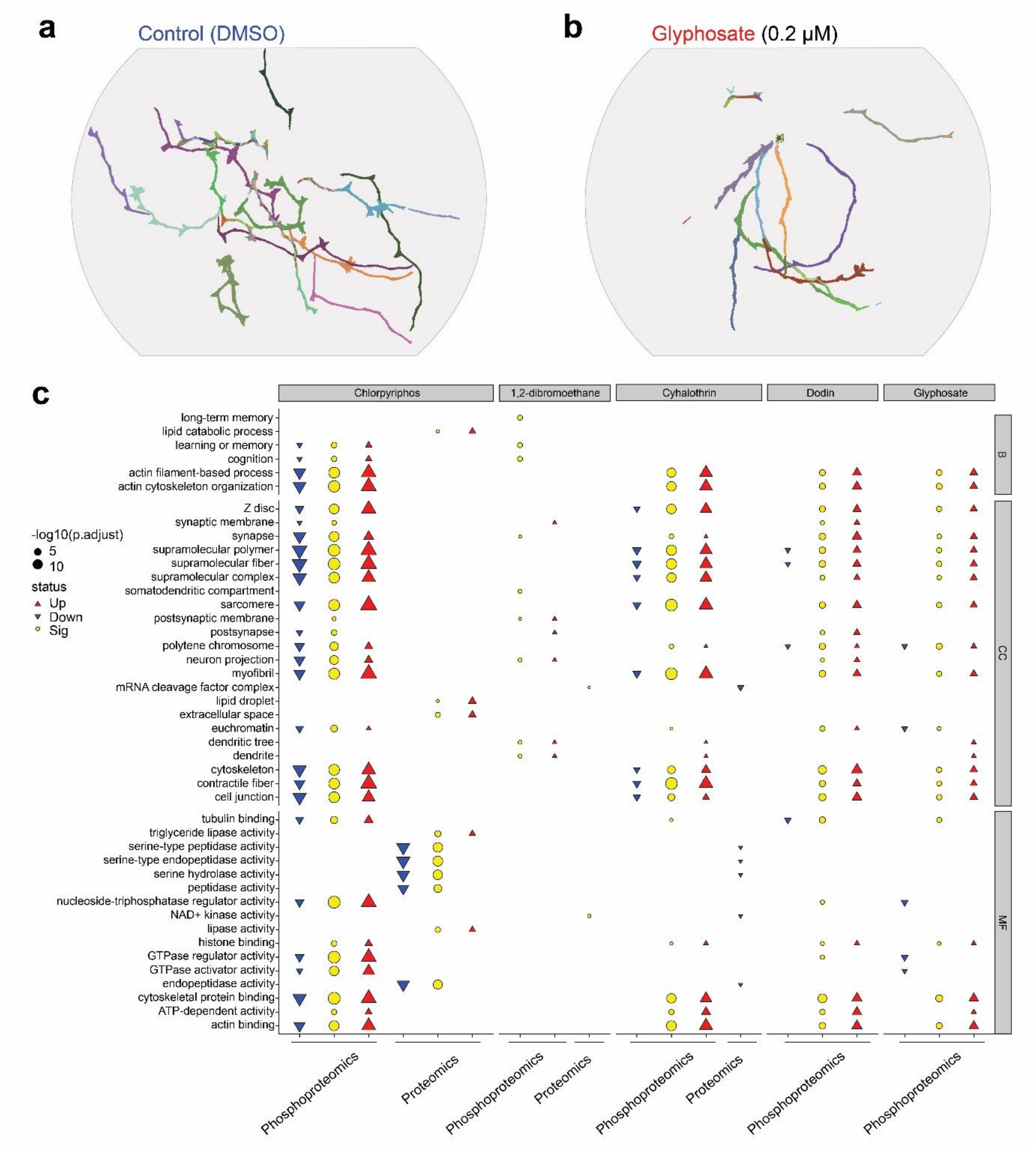


**a-b**, Trajectories of control (**a**) or glyphosate-treated (**b**) larval populations. Each line represents the path of a different larva. The trajectories were obtained using FIMtrack. **c**, Gene Ontology enrichment analysis of hits associated with each treatment (top row), either at the proteomics or phosphoproteomics level (bottom row). Upregulated categories are shown in red, and downregulated ones in blue.

**Extended Data Fig. 4. Temperature-dependent effects of agrochemicals on lethality and behavior.**

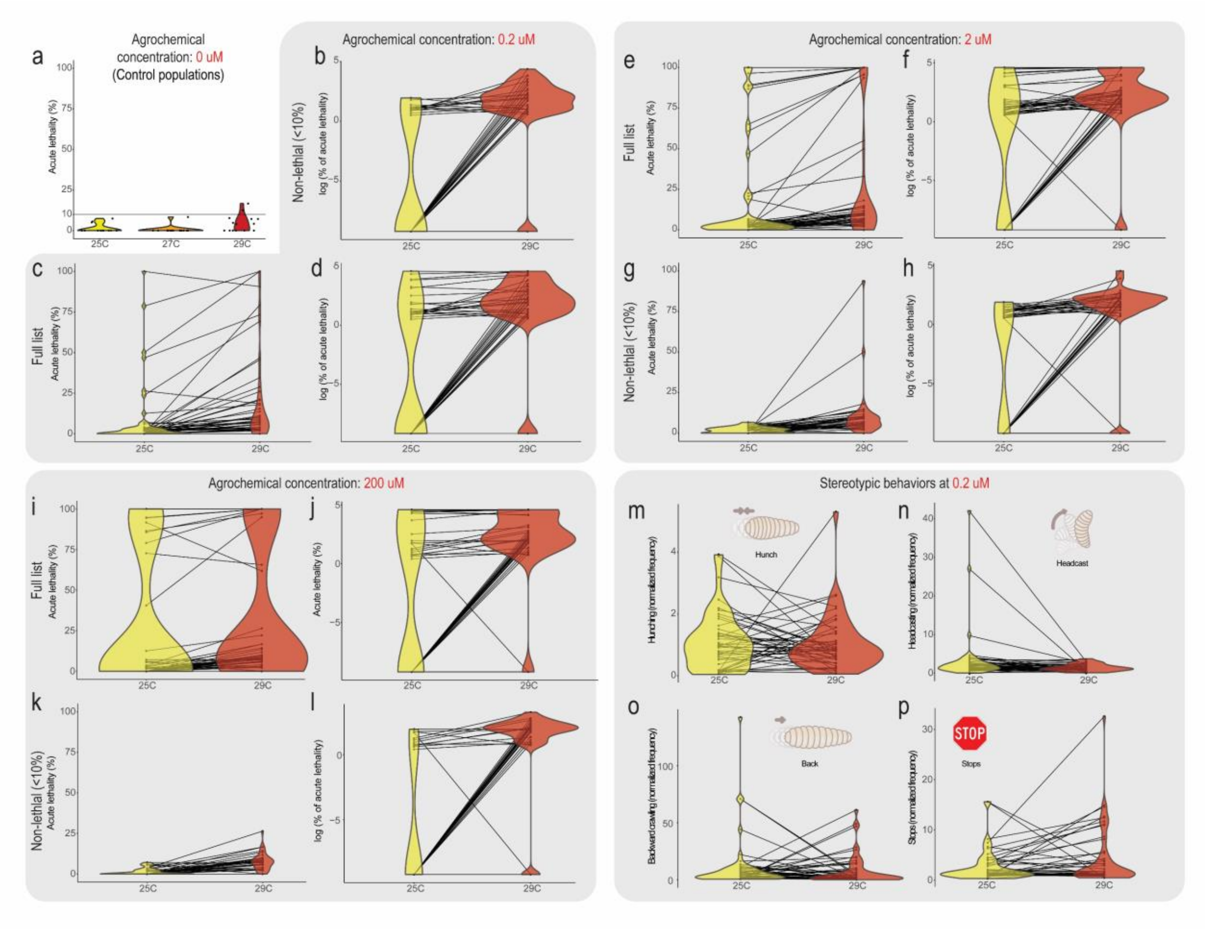


**a,** Average acute lethality (N=2, n~20) observed for all control populations at the reported temperature. The horizontal line indicates the 10% acute lethality threshold used for considering a condition as sublethal. **b-d**, Average acute lethality (N=2, n~20) for pesticides considered non-lethal (<10%) at 25˚C (**b**), or for all 49 agrochemicals used in these assays (**c-d**) at 0.2 µM, either in linear (**c**) or log scales (**b & d**). **e-h**, Average acute lethality (N=2, n~20) for pesticides considered non-lethal (<10%) at 25˚C (**g & h**), or for all 49 agrochemicals used in these assays (**e & f**) at 2 µM, either in linear (**e & g**) or log scales (**f & h**). **i-l**, Average acute lethality (N=2, n~20) for pesticides considered non-lethal (<10%) at 25˚C (**k & l**), or for all 49 agrochemicals used in these assays (**i & j**) at 200 µM, either in linear (**i & k**) or log scales (**j & l**). **m-p**, Average frequencies (N=2, n~20) of different stereotypic movements (hunching in **m**, headcasting in **n**, backward crawling in **o**, and stops in **p**) in populations treated with the 49 agrochemicals used in these assays. Frequencies are normalized to measurements from control populations at the indicated temperature.

**Extended Data Fig. 5. Synergistic interactions between specific agrochemicals.**

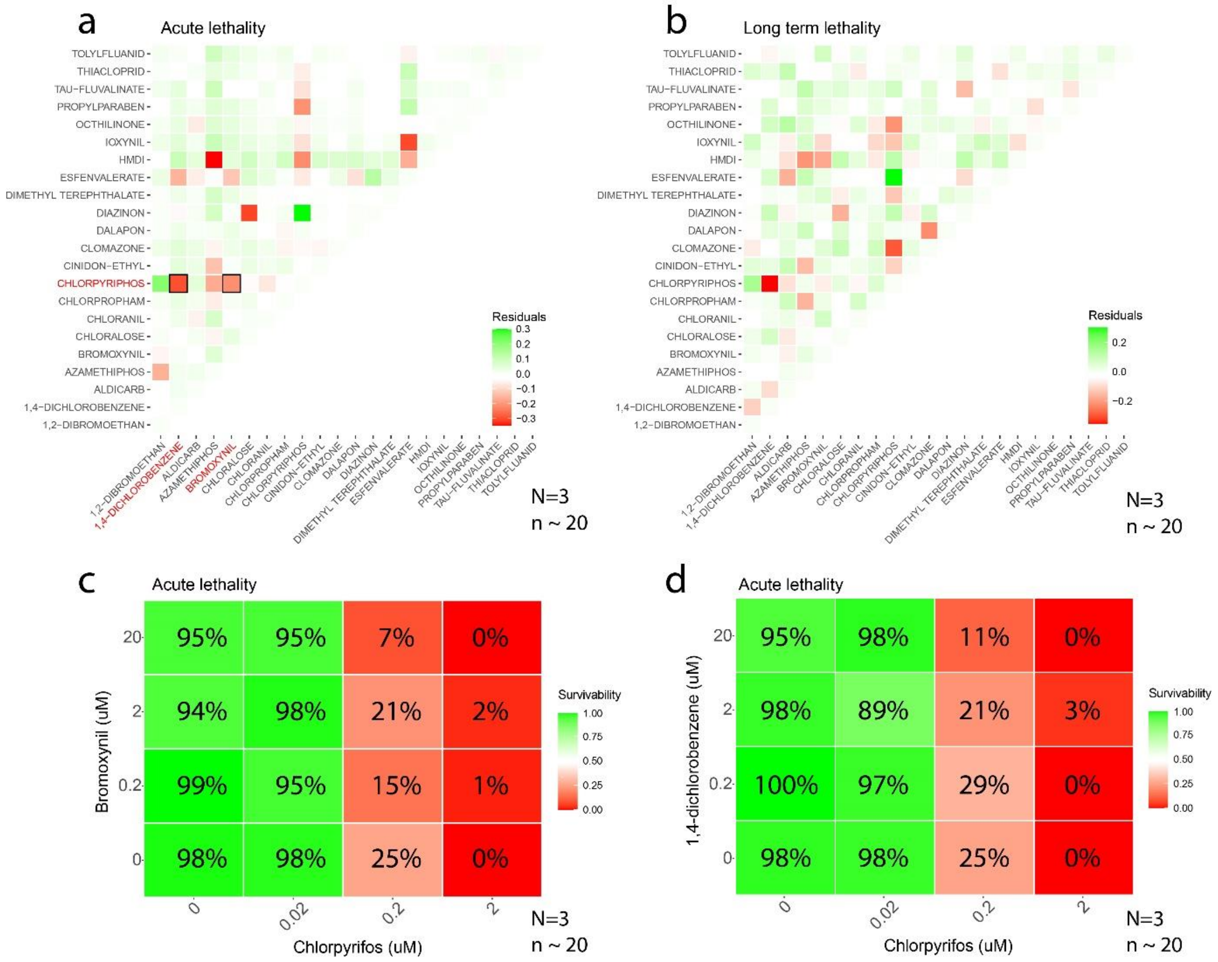


**a-b**, Heat maps showing the difference between observed and expected acute (**a**) or long-term lethality (**b**), assuming a linear interaction between the combined molecules. Green squares (observed>expected) reflect potential antagonistic interactions, while the red ones (observed<expected) reflect potential synergistic interactions. Molecules highlighted in red in (**a**) were selected for testing synergy through checkerboard analyses. **c-d**, Checkerboard analyses showing the average fraction of surviving larvae (survivability) after a 16 h exposure to a combination of chlorpyrifos and bromoxynil (**c**) or 1,4-dichlorobenzene (**d**) at the indicated concentrations. The average percentage of surviving larvae is also shown for each condition.

**Extended Data Fig. 6. Behavioral repertoire of larvae exposed to a combination of pesticides.**

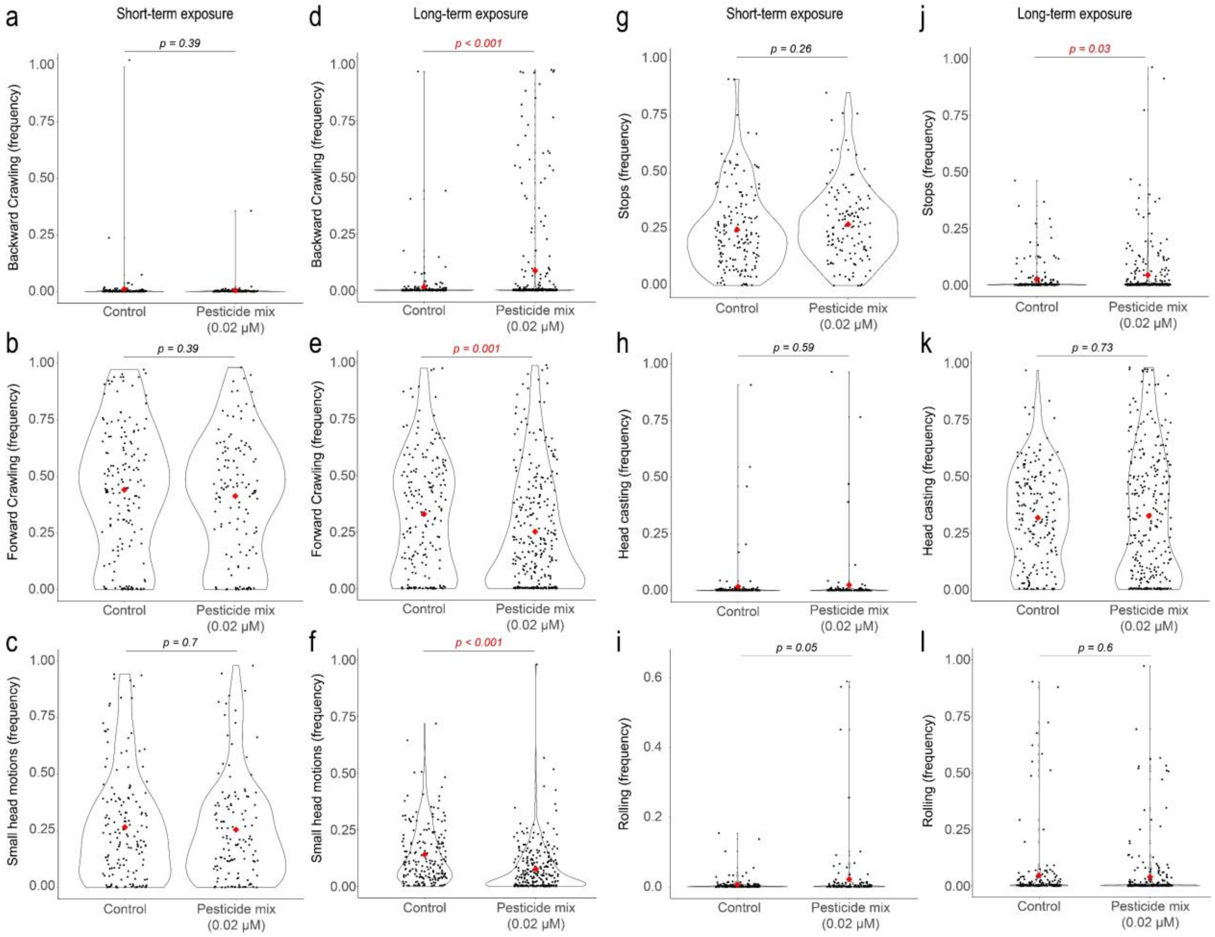


Frequencies of stereotypic behaviors measured in populations exposed to the pesticide mix described in Fig 4b for 16 h (short-term) or 5 days (long-term). Each dot represents an individual larva. Red dots show mean values for each condition across all larvae. p-values for two-tailed T-tests. The significant ones ($p<0.05$) are highlighted in red. N=2, n~20

**Extended Data Fig. 7. Natural isolates show higher resistance to organophosphates.**

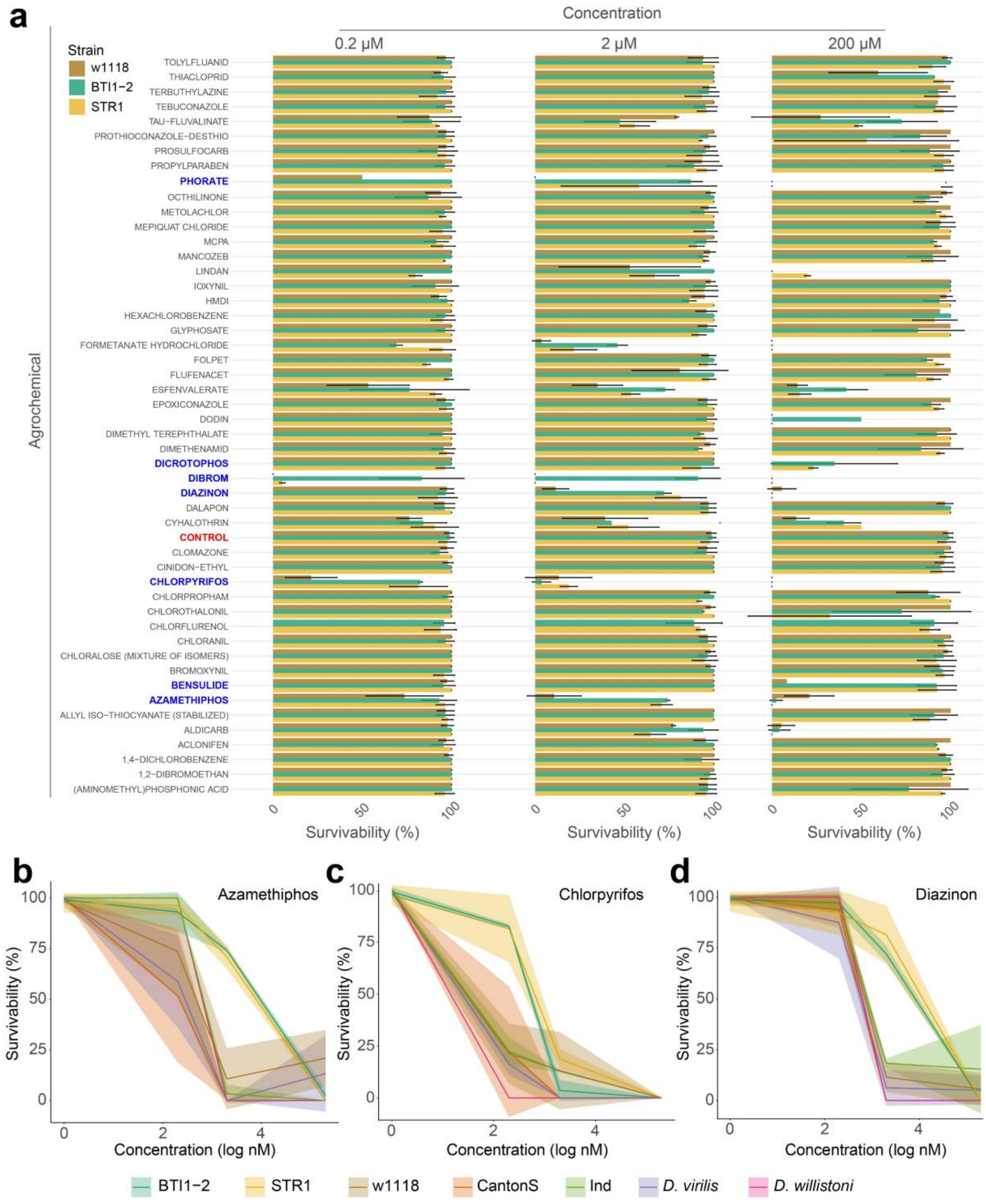


**a**, Percentage of surviving larvae of the indicated strains (color code) after a 16 h exposure to the specified molecules (rows) and concentrations (columns). Control populations are highlighted in red. Organophosphate insecticides are highlighted in blue. **b-d**, Dose-response curves showing the percentage of surviving larvae for different strains (colors) exposed to increasing concentrations of azamethiphos (**b**), chlorpyrifos (**c**) or diazinon (**d**) for 16 h.
